# Modeling Tick Populations: An Ecological Test Case for Gradient Boosted Trees

**DOI:** 10.1101/2023.03.13.532443

**Authors:** William Manley, Tam Tran, Melissa Prusinski, Dustin Brisson

## Abstract

General linear models have been the foundational statistical framework used to discover the ecological processes that explain the distribution and abundance of natural populations. Analyses of the rapidly expanding cache of environmental and ecological data, however, require advanced statistical methods to contend with complexities inherent to extremely large natural data sets. Modern machine learning frameworks such as gradient boosted trees efficiently identify complex ecological relationships in massive data sets, which are expected to result in accurate predictions of the distribution and abundance of organisms in nature. However, rigorous assessments of the theoretical advantages of these methodologies on natural data sets are rare. Here we compare the abilities of gradient boosted and linear models to identify environmental features that explain observed variations in the distribution and abundance of blacklegged tick (*Ixodes scapularis*) populations in a data set collected across New York State over a ten-year period. The gradient boosted and linear models use similar environmental features to explain tick demography, although the gradient boosted models found non-linear relationships and interactions that are difficult to anticipate and often impractical to identify with a linear modeling framework. Further, the gradient boosted models predicted the distribution and abundance of ticks in years and areas beyond the training data with much greater accuracy than their linear model counterparts. The flexible gradient boosting framework also permitted additional model types that provide practical advantages for tick surveillance and public health. The results highlight the potential of gradient boosted models to discover novel ecological phenomena affecting pathogen demography and as a powerful public health tool to mitigate disease risks.

## Introduction

Statistical models have been a cornerstone of understanding ecological phenomena in the natural world. Ecological models traditionally focus on identifying the biotic and abiotic drivers of natural phenomena and on explaining the distribution and abundance of populations (Austin et al., 1984; Elith and Leathwick, 2009; Harvey et al., 1980; McLain et al., 1995; Tran et al., 2021a). Classical generalized linear modeling has resulted in many foundational ecological discoveries (Abbott et al., 1977; Austin et al., 1990; Elith and Leathwick, 2009; Kleiber, 1947; Root, 1988; Tilman et al., 1996). This modeling framework, however, has several technical disadvantages including strict assumptions about error distributions, sensitivity to outliers, and an assumption of linear relationships between variables that can limit predictive power (Hastie et al., 2001; McCullagh and Nelder, 1989; Naghibi and Pourghasemi, 2015; Olden et al., 2008; Yee and Mitchell, 1991). The introduction of machine learning methods such as gradient boosted trees overcomes many of these limitations, although direct comparisons of the effectiveness of machine learning methods and linear models on natural data sets are rare (De’ath, 2007; Elith et al., 2008; Elith et al., 2006; Friedman, 2001). In this study, we compare a gradient boosting machine learning method (Pedregosa et al., 2011) with comparable general linear models in their ability to identify environmental features affecting population dynamics and their ability to predict the distribution and abundance of blacklegged ticks (*Ixodes scapularis*), an arthropod vector of multiple human pathogens.

Many machine-learning frameworks such as neural networks, random forests, and gradient boosted trees are well suited to investigate ecological phenomena in the increasingly data-rich research environment (Cutler et al., 2007; Farley et al., 2018; Friedman, 2001; Han et al., 2015; Rammer and Seidl, 2019; Stephens et al., 2017; Tran et al., 2021b). Among machine learning methods gradient boosted trees are well reputed for very high predictive accuracy and accurate identification of nonlinear relationships on tabular data (Bentéjac et al., 2021; Elith et al., 2008; Grinsztajn et al., 2022). Gradient boosting is an efficient machine learning algorithm that can analyze large data sets, identify complex relationships among variables, and make highly accurate spatio-temporal forecasts. The power of the gradient boosting algorithm is in part derived from their ability to automatically identify non-linear and non-additive relationships by combining hundreds of decision trees into a highly accurate ensemble (De’ath, 2007; De’ath and Fabricius, 2000). These models have several advantages over traditional linear models including that they accept many data types, are unconstrained by data and error distributions, and automatically detect nonlinear and interactive relationships. Further, cross-validation and advances in interpretative machine learning algorithms have addressed prior concerns that gradient boosted algorithms are prone to over-fitting and are too complex to derive ecological inferences (Elith et al., 2008; Lundberg and Lee, 2017; Rudin, 2019; Ryo et al., 2021).

The ability of linear and gradient boosted models to identify ecologically relevant features or to forecast demographic changes is rarely assessed in natural systems, despite the availability of appropriate data sets (though see Becker et al., 2020; Elith et al., 2006; Escobar et al., 2018; Qiao et al., 2015; Shabani et al., 2016). On one such dataset, linear models that explored the explanatory power of 217 environmental variables on the distribution and abundance of *I. scapularis* ticks identified several geographical, temporal, seasonal, environmental, climatic, and landscape features that accounted for the majority of the natural variance in tick demography (Tran et al., 2021a). These linear models accurately predicted the distribution and abundance of tick populations in future years, providing a potentially powerful public health tool to mitigate human disease risks from *I. scapularis*-borne pathogens including the agents causing Lyme disease, babesiosis, and anaplasmosis (Burgdorfer et al., 1982; Spielman et al., 1979; Telford et al., 1996). However, the data distributions assumed in this linear model framework required separate distribution and abundance models and the default assumptions of linearity and additivity limited the exploration of non-linear and non-additive effects which are ubiquitous in ecological systems (Hastie et al., 2001; Levin, 1998; McCullagh and Nelder, 1989; Olden et al., 2008; Tran et al., 2021a; Yee and Mitchell, 1991).

Here, we use gradient boosted trees to investigate the relationship between environmental features and the distribution and abundance of *I. scapularis* using the same dataset previously analyzed with general linear models (Tran et al., 2021a). The gradient boosted models were used to forecast the distribution and abundance of ticks in areas and years not used to build the models. Both the environmental features determined to influence tick demographics and the predictive performance of the gradient boosted tree models were compared to linear models trained and validated using the same data sets (Tran et al., 2021a). Additionally, we utilize the flexibility of the gradient boosting framework to build and validate two additional models that offer practical benefits for disease surveillance, including ease of interpretation and the ability to simultaneously predict tick distribution and abundance.

## Methods

### Study system

The presence and abundance of host-seeking nymphs were determined at 532 unique locations between 2008 and 2018 using the standardized dragging, flagging, and walking survey protocols described previously (Prusinski et al., 2014; Tran et al., 2021a). Locations were sampled every 1–5 years with an average of 4.7 visits per site between 2008 and 2018. The environmental features investigated as explanatory factors in our statistical models can be broadly categorized as geographical, temporal, seasonal, climatic, and landscape features. The tick density and environmental data used in this study are identical to those previously described (Tran et al., 2021a) to rigorously evaluate the relative efficacy of the gradient boosted and linear statistical models.

### Distribution and Abundance Models

Independent distribution and abundance gradient boosted models were built to allow direct comparisons with the previously published distribution and abundance linear models (Tran et al., 2021a). A combined distribution and abundance linear model was not built, as a log-transformation of tick abundance was used to approximate a normal distribution and thus sites where ticks were absent could not be accommodated (Tran et al., 2021a). Data were also processed as described previously (Tran et al., 2021a) to aid comparisons between gradient boosted and linear models. As examples, ticks were considered “present” at a site in a given year if nymphs were detected at any of the multiple site visits within the year and the visit with the greatest nymphal abundance estimate was used as the abundance value for that site in that year. For a summary of built models see (Supplemental Table 2).

Training of gradient boosted models included feature selection, hyper-parameter tuning, and model fitting to the training data set (data from 2008-2017). Environmental features were selected separately for each model using a step-forward feature selection algorithm that optimizes average predictive performance on a 5-fold cross-validation data set (Raschka, 2018). Briefly, each of the 5 folds of the cross-validation data set was generated by randomly partitioning the training data into subsets for model fitting (80% of data) and evaluation (20% of data), such that each fold would contain a unique 20% of the training data for evaluation. Models were limited to 30 or fewer environmental features to reduce the probability of over-fitting (Cawley and Talbot, 2010). Hyper-parameters that influence the learning process were tuned using a random search algorithm to find values that maximized performance on cross-validation data sets (Pedregosa et al., 2011). Using cross-validation sets to optimize which features and hyper-parameters are used in the final model fitting process reduces over-fitting to the training data, making the resultant model more likely to generalize to out-of-sample data (data collected in 2018, which was not used to train the model). The analytical code for this training process is available at MendeleyData (doi: https://doi.org/10.17632/w8bp678m3f.2).

### Predictive Accuracy Assessment

The out-of-sample predictive accuracy of the gradient boosted distribution and abundance models was compared to the accuracy of linear distribution and the abundance models using the previously published accuracy metrics (Tran et al., 2021a). Briefly, the predictions from gradient boosted and linear distribution models to the 2018 out-of-sample data were assessed based on accuracy, sensitivity, and specificity. Abundance model predictions to the out-of-sample data were compared using root-mean-squared-error and R^2^ values. Additionally, to compare the abundance models in accordance with the methodology from (Tran et al., 2021a), abundances were converted from log-transformed counts of nymphs into discrete categories of low (1-4 nymphs), medium (7-35), and high (36+), and predictions were considered accurate if they were within one natural log unit of the average prediction error.

### Simultaneous Modeling of Distribution and Abundance

A multi-class categorical model and a density-estimating regression model were built using the gradient boosting framework. These models do not require the data processing, such as the log-transformation necessary for the linear models, which allows simultaneous analysis of presence and abundance from all sites and years. The multi-class model predicts nymphal abundance to one of three categories: absent (no nymphs), low abundance (1-35 nymphs), and high abundance (*>*35 nymphs). Out-of-sample performance was assessed as the accuracy of the predicted classification to locations visited in 2018.

The gradient boosted density model is similar to the previously described abundance model except that the response variable was tick density, as opposed to the number of ticks collected used in the linear model, and that site densities of zero ticks were permitted. Nymphal density was estimated as the number of ticks collected per collection-hour. Collector hours here were limited to four as preliminary analyses and prior studies demonstrated that density estimates were biased when larger collection-hour values were included (Tran et al., 2021a). The statistical weight of sites during model fitting was positively correlated with collection-hour up to four hours as density estimate accuracy is greater at sites with more sampling effort.

### Environmental Feature Analyses

The relationships between nymphal tick distribution or abundance with individual environmental features in each model were analyzed using SHAP (SHapley Additive exPlanation) values (Lundberg and Lee, 2017). Briefly, this interpretative framework estimates the impact each model feature has on model predictions. Together these estimates provide a global view of the impact of each feature on model predictions in the context of other model features. SHAP values were used to identify and visualize the non-linear relationships and interaction effects discovered by each model. SHAP values were not used to evaluate the impact of environmental variables on predictions from the multi-class model as the complex outputs of this model are not supported in this analytical framework.

## Results

The gradient boosted distribution and abundance models outperformed their linear model counterparts in both predictive power and identification of complex relationships between environmental features. The gradient boosted distribution model (Figure 1A), built using data from 2008-2017, accurately predicted 94% of sites where ticks were present in 2018 and 84% of sites where ticks were absent. By comparison, the linear distribution model trained and tested on the same data accurately predicted 80.6% of sites where ticks were present and 80.7% of sites where they were absent. Importantly, the gradient boosted model had a far lower false negative rate than the linear model (5.8% vs 19.4%), an especially costly error for public health efforts. The gradient boosted distribution model also made highly accurate predictions to the 27 sites that were visited for the first time in 2018 (true positive rate = 85%; true negative rate = 86%).

**Figure 1:**
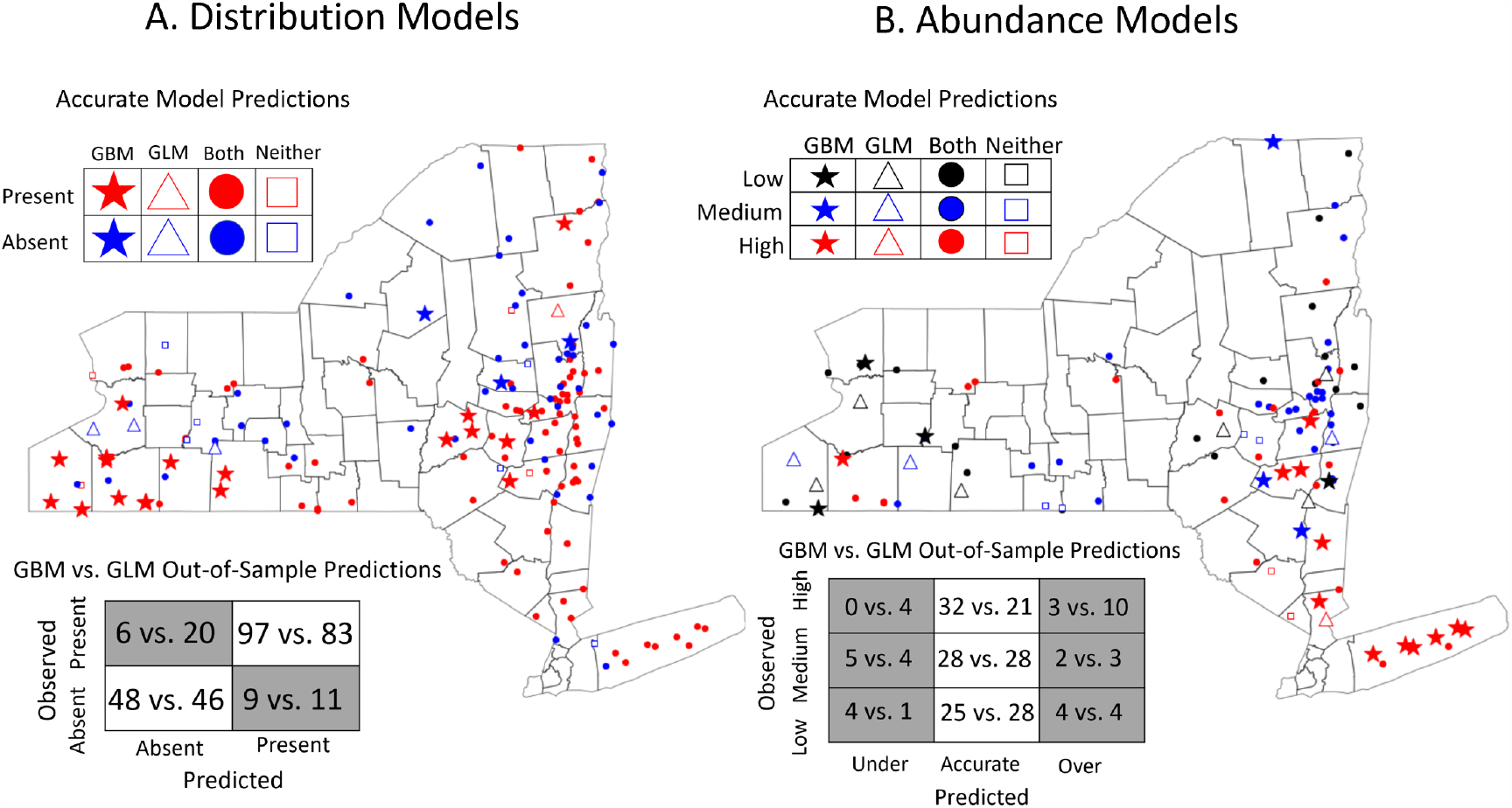
Gradient boosted models more accurately predict future (A) distributions and (B) abundances of nymphal ticks than generalized linear models. (A) The gradient boosted distribution model was more accurate (90.6% vs 80.6%), more sensitive (true positive rate = 94.2% vs 80.5%), and more specific (true negative rate = 84.2% vs 80.7%) than its linear model analog. (B) The gradient boosted abundance model also more accurately predicted to the out-of-sample data than its linear model counterpart (82.5% vs 74.8%). Stars indicate sites with accurate predictions from the gradient boosted model and inaccurate predictions from the linear model; triangles represent accurate linear model predictions and inaccurate gradient boosted model predictions; squares represent sites accurately predicted by both models; circles represent inaccurate predictions by both models. Confusion matrices summarize the accurate and inaccurate predictions made by the gradient boosted model vs the linear model.

The gradient boosted abundance model more accurately predicted out-of-sample tick abundance than the analogous linear model in all quantitative metrics (RMSE = 0.972 vs. 1.096; R^2^ = 0.59 vs. 0.48). Gradient boosted model predictions were also converted into discrete categories to compare the accuracy of the linear and gradient boosted models using the previously published methodology (Tran et al., 2021a). The gradient boosted abundance model was more accurate than its linear model counterpart, correctly predicting the abundance at 82.5% of sites compared to the 74.8% of sites correctly predicted by the linear model (Figure 1B). Sites visited for the first time in 2018 were also predicted with high accuracy by the gradient boosted model (83.3%; RMSE = 0.948; R^2^ = 0.61). Importantly, nearly 40% of all sites incorrectly predicted by the gradient boosted model were conservative in that the model overestimated tick abundances at sites with high abundance (n=3) or underestimated tick abundance at sites with low abundance (n=4). These errors are less costly as they indicate that the model has correctly predicted sites with high or low tick abundance but erred in terms of magnitude.

Complex non-linear relationships between environmental features and nymphal abundance were detected in gradient boosted models that were not investigated in the previously published linear models (Tran et al., 2021a). For example, estimates of deer population size have a highly complex relationship with nymphal abundance (Figure 2A): deer harvest values less than 2000 result in decreased nymphal abundance predictions; deer harvest between 2000 and 3000 are correlated with increases in nymphal abundances; deer harvest between 3000 and 6000 are correlated with decreased nymphal abundances; and deer harvest above 6000 is correlated with increased nymphal abundance. Although not biologically relevant, the number of tick collection efforts (sampling hours) had a positive but decelerating relationship with the number of nymphs collected (Figure 2B). That is, the number of nymphs collected is strongly and positively correlated with the number of hours field technicians flagged for ticks at sites visited for fewer than two hours. However, this positive relationship becomes less pronounced at sites visited for greater than two hours and is not detectable at sites visited for more than five hours.

**Figure 2:**
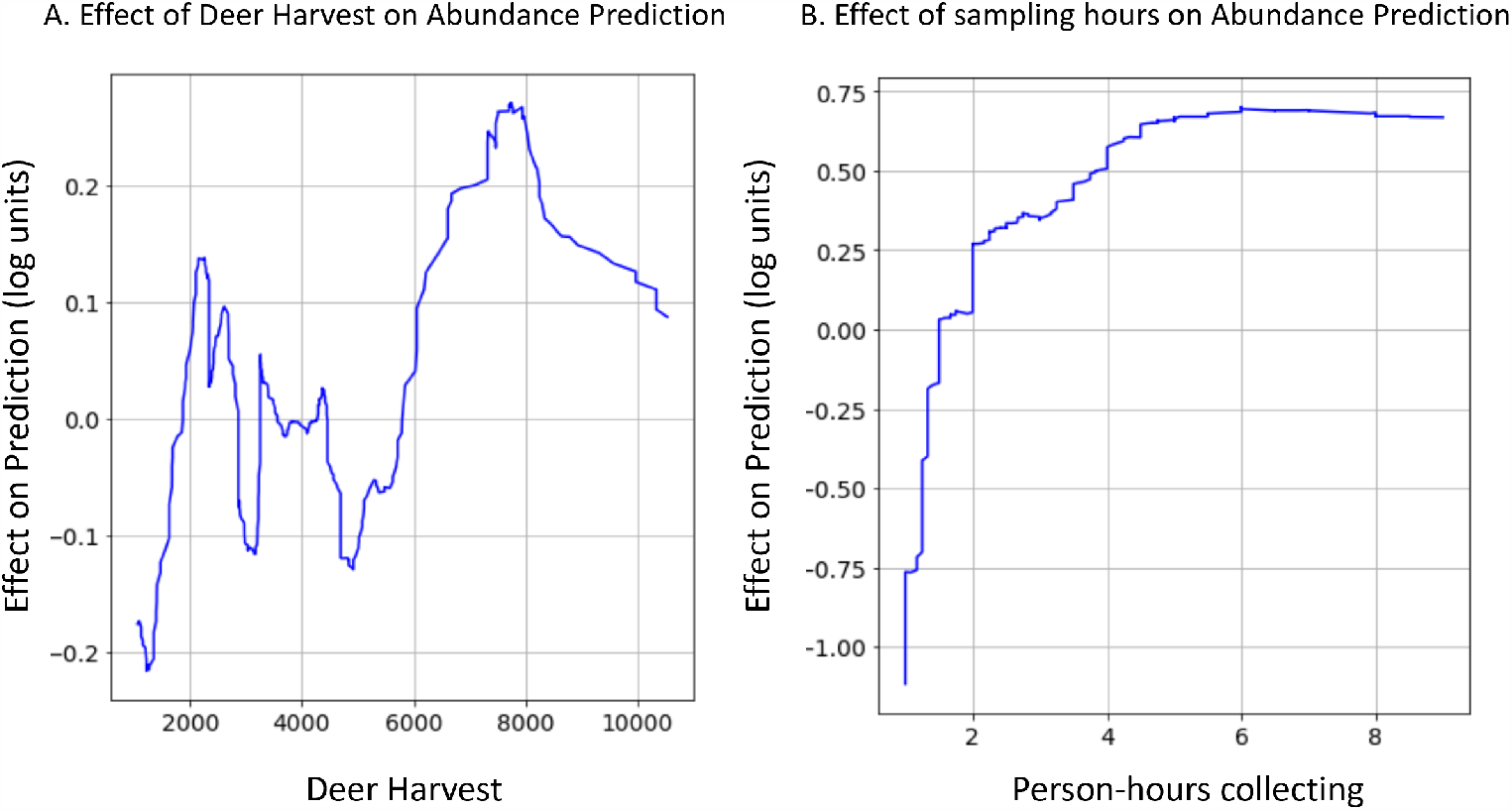
Gradient boosted models identified non-linear relationships that are impractical to investigate with linear models. (A) The association between estimates of deer population size and nymphal tick abundance oscillates between having a positive effect to a negative effect. (B) The relationship between person-hours collecting hours and tick abundance is a positive but decelerating function. Data shown are the rolling average (rolling window = 50) of the impact that (A) deer density estimates or (B) tick collection effort has on tick abundance.

The impacts of non-additive interactions between environmental features on the presence of nymphal ticks were also detected in gradient boosted models. One ecologically relevant interaction demonstrates that the effect of the month in which a site is sampled on the presence of active nymphs is conditioned on the maximum temperature in June of the year before sampling (Figure 3). Although sampling month is generally highly predictive of nymphal presence due to the seasonal activity patterns of *I. scapularis* in New York State (Yuval and Spielman, 1990), ticks were more likely to be detected in the summer months (May-August) if the temperature in June of the prior year was hotter. By contrast, the probability of detecting nymphal ticks in fall months (September-December) was greater if the maximum temperature in June of the prior year was cooler. This non-additive effect was strong enough to change the month of May from being negatively associated with the presence of nymphs when June of the prior year was cooler to a positive association when this month was warmer.

**Figure 3:**
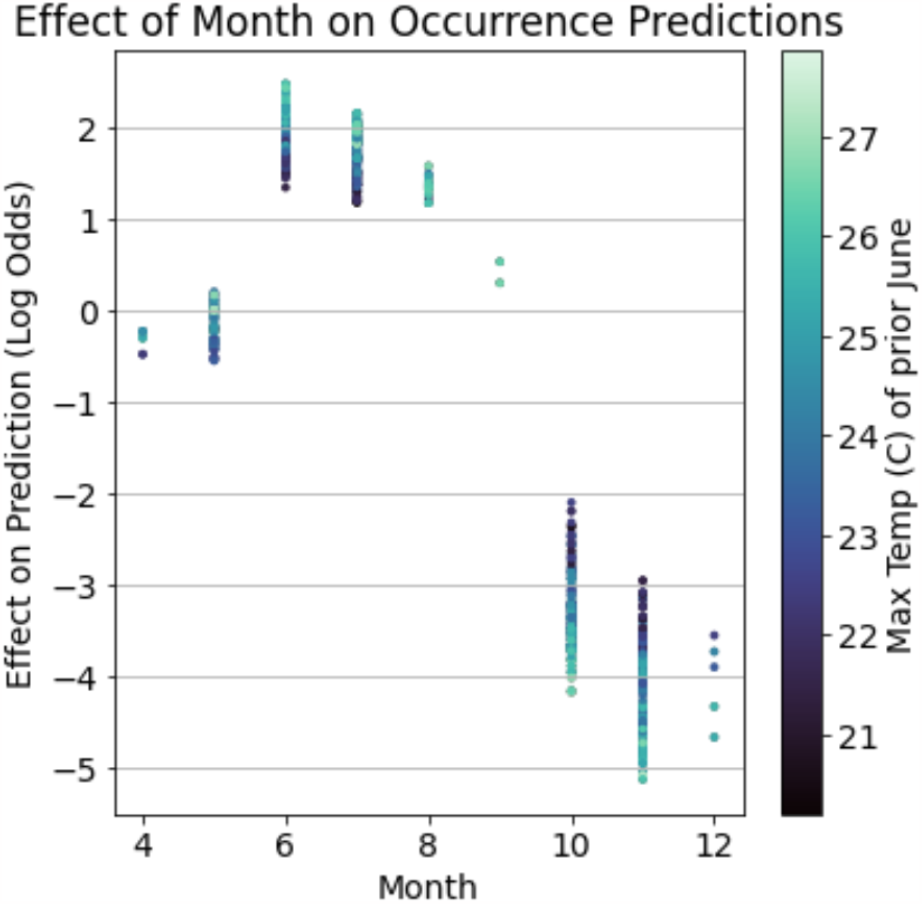
Gradient boosted models detected ecologically relevant interactions between environmental features which impacts the presence of nymphal ticks. The maximum temperature in June of the year before a collection event modulates tick phenology. That is, nymphal ticks are more likely to be collected between May and August in years when the prior June was hotter while the likelihood of nymphal tick presence in September-December increases in years when the prior June was cooler.

The sets of environmental features used by the gradient boosted distribution and abundance models were similar to those included in linear models but were related to nymph populations in more complex ways. Despite different feature selection processes, the two modeling frameworks frequently used identical or strongly correlated features as predictors (Supplement Table 1). However, the linear models related features to nymph populations linearly and without interaction effects, while the relationships in the gradient boosted models were always non-linear and frequently incorporated interactions. In fact, both non-linear relationships discussed above (Figure 2) involve features that were included in the previously published linear models.

The gradient boosting framework was used to produce two additional models - a multi-class and a density model - that simultaneously estimate the presence and abundance of nymphs. The multi-class model forecasts which sites will have no nymphs, low nymphal abundance (1-35), or high nymphal abundance (*>*35) with high accuracy, correctly classifying 80% of sites in the out-of-sample data set (Figure 4). This multi-class model predicted the presence or absence of nymphs with similar accuracy as the gradient boosted distribution model (both ≈90%) but has the additional functionality of distinguishing between two non-zero abundance classes. The novel density model predicts a continuous estimate of tick densities (ticks per collection hour) to out-of-sample data with high accuracy (R^2^ = 0.42). Restricting the comparison to the subset of the out-of-sample data included in the abundance models (Figure 1B) resulted in the density model performing comparably with the linear abundance model (RMSE = 1.06 vs. 1.096; R^2^ = 0.51 vs. 0.48) while retaining the added functionality of predicting the absence of nymphs. Both the multi-class and density models have similar predictive accuracy at sites that were visited for the first time in 2018 and those that had been sampled prior to 2018.

**Figure 4:**
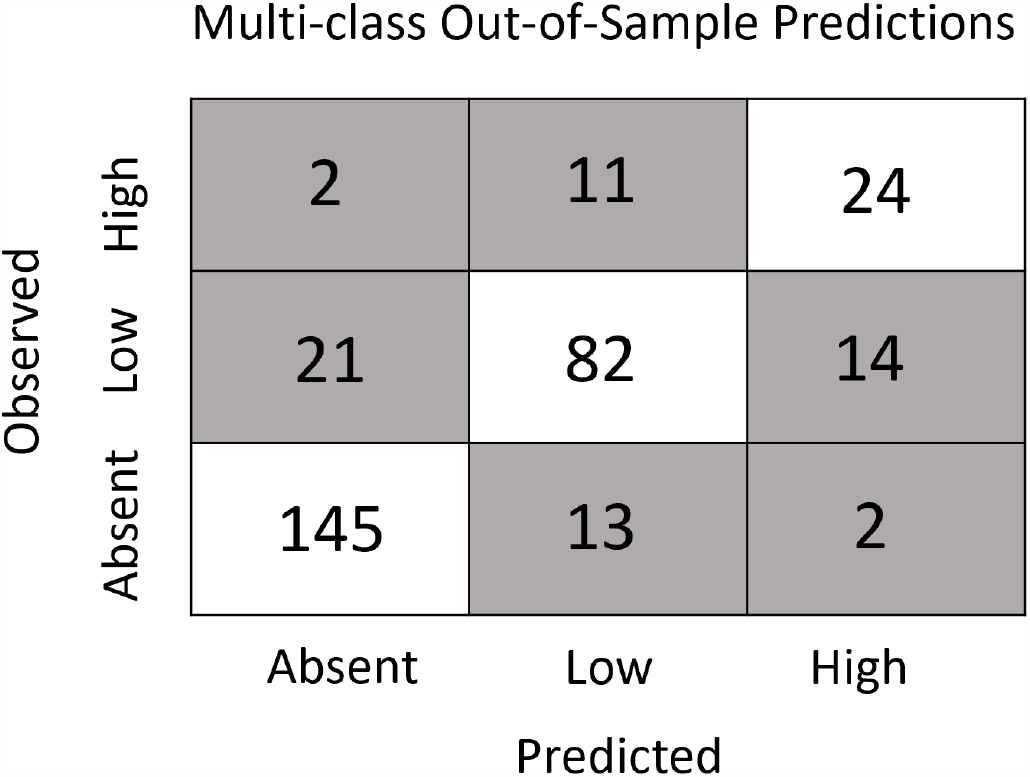
The multi-class model accurately predicts both the presence and abundance of nymphs across New York State. The model accurately predicted 90.6% of sites without ticks, 70% of sites with low tick abundance (1-35), and 64.9% of sites with high tick abundance (*>* 35). Further, most inaccurate predictions were one class apart (absent vs low or low vs high). That is, sites without nymphs were rarely predicted to have a high abundance (1.3%) and sites with high abundance were rarely predicted to have no nymphs (5.4%).

## Discussion

Machine learning analyses of the recent expansion of publicly available biological and environmental data is ideal for discovering novel ecological insights and accurately forecasting the distribution and abundance of populations in nature. The gradient boosted modeling framework efficiently and accurately identifies both simple and complex ecological relationships from large data sets and produces highly accurate predictions of the demography of natural populations (Elith et al., 2008; Han et al., 2015; Ramazi et al., 2021; Wyse and Dickie, 2018). However, the theoretical advantages of gradient boosted models over traditional linear models are rarely validated using natural data sets. As a result, many ecologists rely exclusively on generalized linear models even though gradient boosted models could be more effective for exploring and interpreting data (LaRue et al., 2019; Shah et al., 2019; Sutomo et al., 2021; Walter et al., 2018). Here we demonstrate that the distribution and abundance of natural populations of *I. scapularis* ticks can be predicted with greater efficiency and accuracy with gradient boosted models than with linear models. Additionally, the gradient boosted models identified non-linear and non-additive relationships, which are difficult to detect in linear modeling frameworks, that improved predictive accuracy. These results indicate that gradient boosted models can improve both spatio-temporal forecasts and provide novel insights into the ecology of natural populations.

The gradient boosted occurrence and abundance models consistently outperformed their linear counterparts in predictive accuracy, illustrating the potential of this framework to improve predictions of ecological phenomena. When trained and tested on the same datasets as the linear models from (Tran et al., 2021a), the gradient boosted models were better able to forecast the distribution and abundance of nymphs (Figure 1). Notably, the gradient boosted models outperformed their linear analogs on sites not previously sampled, suggesting that the superior predictive performance of this framework results from incorporating more precise ecological relationships rather than overfitting to previously sampled sites. However, gradient boosted models are not always expected to be the most accurate type of model for a given problem. As examples, linear models might be favored for small datasets with simpler relationships when overfitting is likely to be a problem, whereas neural networks are expected to outperform in contexts like image or speech classification (Deng et al., 2013; Hastie et al., 2001; Rawat and Wang, 2017). Nonetheless, our findings highlight gradient boosted models as a powerful but underutilized tool for predicting demographic changes in natural populations.

The gradient boosted models automatically identified complex relationships between several environmental features and the distribution and abundance of ticks. For example, these models found a non-linear relationship between deer harvest data - an estimate of deer population size - and nymphal tick abundance (Tran et al., 2021a). The non-linear relationship identified in the gradient boosted model implies that changes in deer populations are positively associated with tick abundance at some deer population sizes and negatively at others (Figure 2). This non-linear relationship may explain contradictory conclusions in previous reports in which some identify positive relationships between deer population size and tick densities while others do not (Kugeler et al., 2016; Lewis et al., 2017; Ostfeld et al., 2006; Schulze et al., 2001; Tran et al., 2021a). Statistical models like gradient boosting do not identify the ecological mechanism underlying this relationship but do suggest avenues for further experimentation to resolve this discrepancy. Gradient boosted models also identified an interaction between climate variables that influences tick questing activity throughout summer months. Specifically, hotter temperatures in June of the year prior to tick collections alter tick phenology such that nymphal ticks are active earlier in the season (Figure 3). These results warrant further investigation into how climate change may affect seasonal activity patterns of ticks and possibly the pathogens they transmit (MacDonald et al., 2021).

Relationships between variables identified by any statistical model should be interpreted with caution. The ecological relationships included in the gradient boosted models presented here were identified using SHAP value analyses that determine the effect each variable has on model predictions (Lundberg and Lee, 2017). Thus, these relationships represent the patterns our models used to make accurate predictions but do not necessarily represent causal processes. Nevertheless, similar environmental features were detected in the gradient boosted and linear models despite using different approaches (Supplemental Table 1), adding confidence that these features are useful in forecasting tick distribution and abundance (Tran et al., 2021a). Additionally, the complex relationships involving these shared environmental features suggests that the gradient boosted framework has the potential to yield novel ecological insights, even on datasets previously analyzed with traditional statistical methods. While further experimentation is needed to clarify the biological significance of these relationships, they demonstrate the ability of the gradient boosting framework to automatically discover non-linear and interaction effects which general linear models often do not detect.

The flexibility of the gradient boosted modeling framework allowed us to build models with at least three practical advantages for both ecological interpretation and public health (De’ath, 2007). First, the multi-class and density model simultaneously predict the distribution and abundance of ticks, allowing tick population size to be estimated with a single model. Second, data pre-processing such as log-transformations is not required in the gradient boosting framework making both the predictions and error estimates more interpretable. Lastly, the density model analyzes tick density directly, a correlate of the human contact risk with a questing nymph, as opposed to the number of ticks collected which is conditioned by the sampling effort (Khatchikian et al., 2012). While it is in principle possible to achieve these advantages using generalized linear models (for an ecological example see Bah et al., 2022), the flexibility of the gradient boosting framework greatly simplified the process of implementing these multiple types of models (Natekin and Knoll, 2013).

Applying the gradient boosted modeling framework to pathogens carried by *I. scapularis* may provide additional improvements for disease risk forecasting and could identify the environmental features that correlate with human risk of contracting a *I*.*scapularis*-borne disease. For example, gradient boosted analyses of the distribution and abundance of ticks carrying *Borrelia burgdorferi, Babesia microti, Anaplasma phagocytophilum*, or other tick-borne pathogens are likely to identify ecological factors impacting pathogen populations and could predict the risk of encountering an infected tick. More broadly, the gradient boosted framework can improve ecological models of many infectious disease systems (Ashby et al., 2017; Fischhoff et al., 2021; Giles et al., 2018; Han et al., 2015; Solano-Villarreal et al., 2019). The rapidly expanding environmental data sets can be efficiently analyzed by gradient boosted models in order to detect ecological relationships and accurately predict disease risk in many systems, thus promoting a better understanding of natural disease systems and aiding the development of public health strategies.

## Data Availability

Data used to train and validate models are from (Tran et al., 2021a). Data and code for model training and evaluation are available at MendeleyData (doi: https://doi.org/10.17632/w8bp678m3f.2).

## Funding

This work was supported by the NYSDOH, the National Institutes of Health (AI142572), and the Burroughs Welcome Fund (1012376).

## Conflict of Interest Disclosure

The authors of this preprint declare that they have no financial conflict of interest with the content of this article. D Brisson is a Recommender at PCI Ecology and is on the Managing Board at PCI Evolutionary Biology.

## Appendix A Supplementary data

**Supplemental Table 1:** Most Predictive Ecological Features from Gradient Boosted Occurrence and Abundance Models compared to Linear Counterparts.

| Model | GBM<br>Occurrence | GLM<br>Occurrence | GBM<br>Abundance | GLM<br>Abundance |
| --- | --- | --- | --- | --- |
| **Physical Habitat** | Longitude (+, NL),<br>Distance to nearest road (-, NL) | Latitude (+),<br>Elevation (-),<br>Distance to nearest road (+),<br>Road type of nearest road (NL),<br>Indicator of critical zone (-) | Latitude (-, NL),<br>Longitude (+, NL) | Latitude (-),<br>Longitude (+),<br>Elevation (NL),<br>Forest (-),<br>Distance to nearest hydrography feature (-) |
| **Vapor Pressure** | Maximum Jan 2 years prior (-, NL),<br>Minimum Oct 2 years prior (NL),<br>Maximum Oct 1 year prior (+, NL),<br>Maximum Jan (-, NL),<br>Minimum June (-, NL),<br>Minimum October (+, NL) | Minimum Jan 1 year prior (-) |  | Maximum October 2 years prior (+),<br>Minimum October 2 years prior (-, NL) |
| **Temperature** | Mean differential Jan 2 years prior (+, NL, IE),<br>Degree days above 0 C spring-summer 1 year prior (+, NL, IE),<br>Degree days above 0 C spring 1 year prior (+, NL),<br>Maximum June 1 year prior (+, NL, IE) | Degree days above 0 C spring 2 years prior (-),<br>Degree Days below 0 C winter 1 year prior (+),<br>Degree days above 0 C spring-summer 1 year prior (+) |  | Degree days above 0 C spring-summer 1 year prior (+) |
| **Day of Collection** | Person-hours collecting (+, NL),<br>Month (NL) | Person-hours collecting (+),<br>Month (NL),<br>Local Temperature (+),<br>Wet (-) | Person-hours collecting (+, NL),<br>Week (NL) | Person-hours collecting (+),<br>Month (NL) |
| **Miscellaneous** |  | Deer harvest (-) | Deer harvest (NL) | Deer harvest (+) |

Top 15 most predictive features from the gradient boosted occurrence model and all features from the other models are included.
(-) = negative relationship, (+) = positive relationship, NL = nonlinear relationship, IE = interaction effect

Supplemental Data 1: Table containing all features used by the gradient boosted models can be found at: MendeleyData (doi: https://doi.org/10.17632/w8bp678m3f.2).

**Supplemental Table 2:** Summary of Model Characteristics.

| **Model** | **Sites Predicted** | **Target Variable** | **Accuracy Metrics for Out of Sample Test** | **GLM Analog** |
| --- | --- | --- | --- | --- |
| **GBM Distribution** | All sites | Binary (Nymphs Present or Absent) | Accuracy, Sensitivity, Specificity | GLM Distribution Model |
| **GBM Abundance** | Sites with Nymphs | Log-transformed Nymph Abundance | RMSE, R<sup>2</sup>, Categorical Accuracy | GLM Abundance Model |
| **GBM Multi-Class** | All sites | Three Abundance Classes of Nymphs | Accuracy | N/A |
| **GBM Density** | All sites | Nymph Abundance/<br>Sampling Hour | RMSE, R<sup>2</sup> | N/A |

Model characteristics of all four gradient boosted models are included.

## Notes

https://doi.org/10.17632/w8bp678m3f.2

### Competing Interest Statement

The authors have declared no competing interest.

### Summary of Updates

To indicate peer review and recommendation by PCI Ecology

